## Supplementary material for "Simple and highly efficient detection of PSD95 using a nanobody and its recombinant minibody derivative": Fig. S1, Fig. S2, Table S1

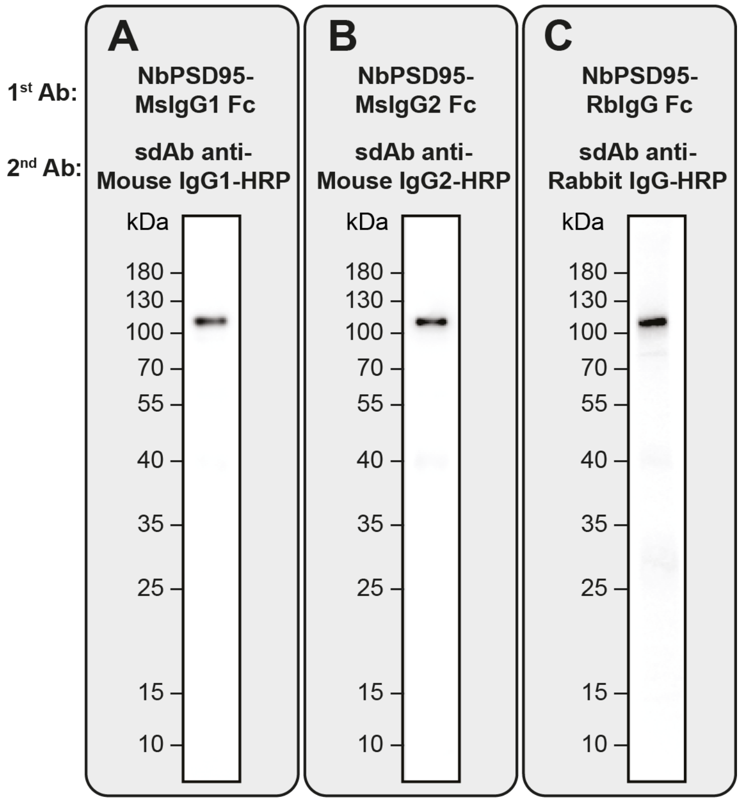


**Figure S1: anti-PSD95 minibodies harboring different Fc domains specifically recognize PSD95 in Western blot applications.**

Total mouse brain lysates were analyzed by Western blot using minibodies consisting of NbPSD95 fused to a Fc domain from either mouse IgG1 (MsIgG1 Fc; **A**), mouse IgG2 (MsIgG2 Fc; **B**) or rabbit IgG (RbIgG Fc; **C**). Detection was performed using HRP-coupled secondary nanobodies recognizing the respective Fc domains. All three minibodies detected PSD95 with similar specificity.

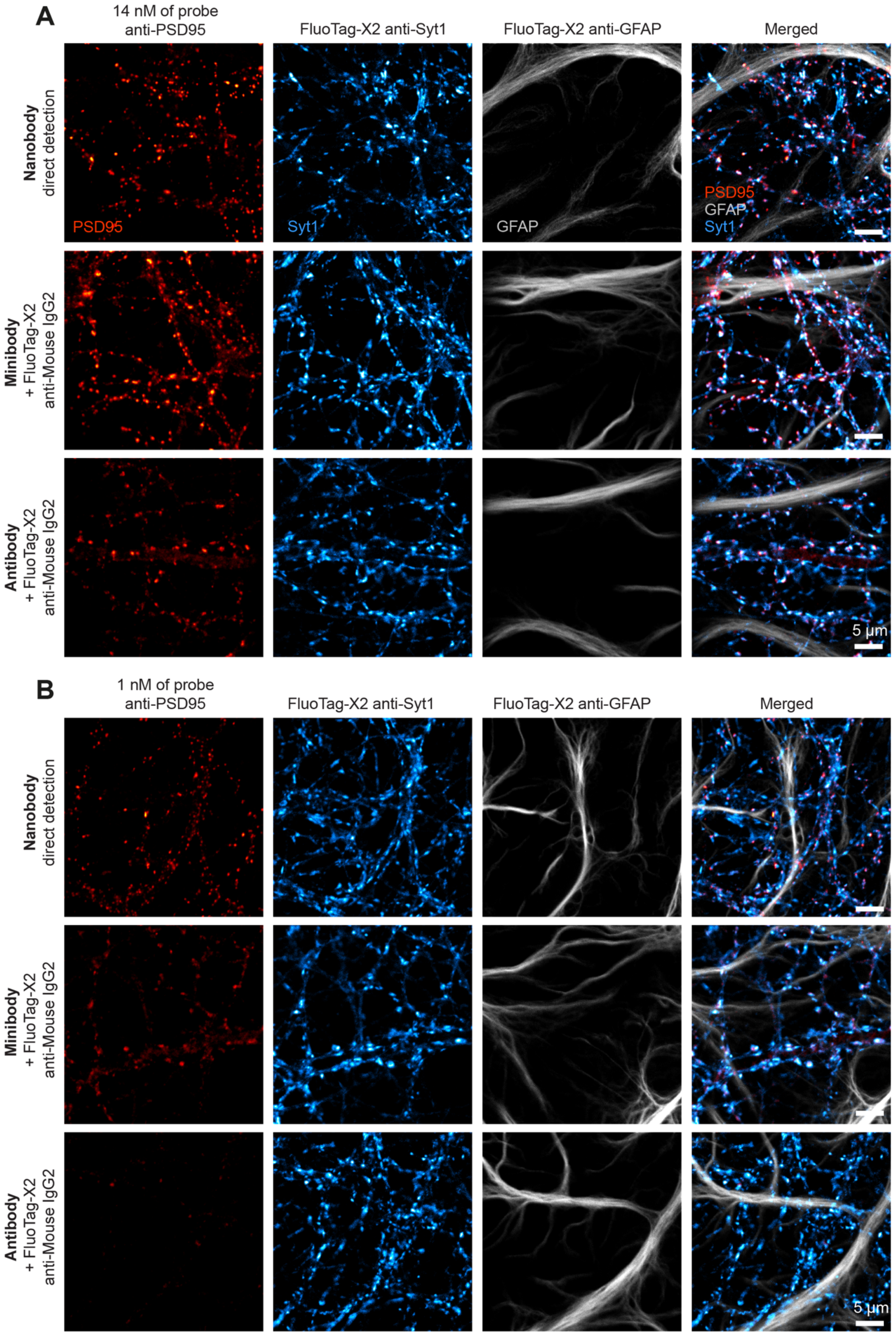


**Figure S2: Comparison of NbPSD95 with an anti-PSD95 minibody and a conventional antibody – related to Fig. 3.** Figure shows separate channels used for creating overlays displayed in Fig. 3.

**Table S1: Antibodies**

| Figure | Panel(s) | Product | Product number | Provider | Description | Application | Dilution/ Conc. |
|---|---|---|---|---|---|---|---|
| Fig. 2A | | Monoclonal anti-FLAG HRP | A8592 | Merck | HRP-conjugated anti-FLAG antibody (clone M2) | ELISA | 1:5000 |
| Fig. 2B | upper panels | FluoTag-X2 anti-PSD95 AZDye568 | N3702-AF568-L | NanoTag Biotechnologies | Fluorophore-conjugated NbPSD95 | IF, direct | 1:500 |
| Fig. 2C | upper left panel | Recombinant anti-PSD95 Antibody (Minibody), Mouse IgG1 Fc fusion | N3782 | NanoTag Biotechnologies | NbPSD95 fused to mouse IgG1 Fc domain | WB, 1st Ab | 1:1000 |
| Fig. 2C | lower panels | Mouse anti-beta-Actin | 251 011 | Synaptic Systems | Monoclonal antibody | WB, 1st Ab | 1:1000 |
| Fig. 2C | upper right panel + lower panels | sdAb anti-MsIgG1 HRP | N2005-HRP | NanoTag Biotechnologies | HRP-conjugated anti-Mouse IgG1 nanobody | WB, 2nd Ab | 1:4000 |
| Fig. 2C | upper right panel | Recombinant anti-PSD95 Antibody (Minibody), Rabbit Fc fusion | N3783 | NanoTag Biotechnologies | NbPSD95 fused to rabbit IgG Fc domain | WB, 1st Ab | 1:1000 |
| Fig. 2C | upper right panel | sdAb anti-RbIgG HRP | N2405-HRP | NanoTag Biotechnologies | HRP-conjugated anti-Rabbit IgG nanobody | WB, 2nd Ab | 1:4000 |
| Fig. 2D | | FluoTag-X2 anti-PSD95 AZDye568 | N3702-AF568-L | NanoTag Biotechnologies | Fluorophore-conjugated NbPSD95 | IF, direct | 1:500 |
| Fig. 2D | | FluoTag-X2 anti-GFAP Atto488 | N3802-At488-L | NanoTag Biotechnologies | Fluorophore-conjugated anti-GFAP nanobody | IF, direct | 1:500 |
| Fig. 3A, Fig. 3B | upper row | FluoTag-X2 anti-PSD95 AbberiorStar635P | N3702-Ab635P-L | NanoTag Biotechnologies | Fluorophore-conjugated NbPSD95 | IF, direct<br>IF, direct | 14 nM<br>1 nM |
| Fig. 3A, Fig. 3B | middle row | Recombinant anti-PSD95 Antibody (Minibody), Mouse IgG2 Fc fusion | N3785 | NanoTag Biotechnologies | NbPSD95 fused to mouse IgG2 Fc domain | IF, 1st Ab<br>IF, 1st Ab | 14 nM<br>1 nM |
| Fig. 3A, Fig. 3B | lower row | Monoclonal anti-PSD95 antibody | MABN68 | Millipore | Monoclonal antibody (Mouse IgG2) | IF, 1st Ab<br>IF, 1st Ab | 14 nM<br>1 nM |
| Fig. 3A, B | two lower rows | FluoTag-X2 anti-Mouse IgG2 AbberiorStar635P | N2702-Ab635P | NanoTag Biotechnologies | Fluorophore-conjugated anti-Mouse IgG2 nanobody | IF. 2nd Ab | 1:500 |
| Fig. 3 | | FluoTag-X2 anti-Syt1 AZDye568 | N2302-AF568-L | NanoTag Biotechnologies | Fluorophore-conjugated anti-Synaptotagmin1 nanobody | IF, direct | 1:500 |
| Fig. 3 | | FluoTag-X2 anti-GFAP Atto488 | N3802-At488-L | NanoTag Biotechnologies | Fluorophore-conjugated anti-GFAP nanobody | IF. direct | 1:500 |

**Table S1: Antibodies (continued)**

| Figure | Panel(s) | Product | Product number | Provider | Description | Application | Dilution/ Conc. |
|---|---|---|---|---|---|---|---|
| Fig. 4 | | FluoTag-X2 anti-PSD95 AbberiorStar635P | N3702-Ab635P-L | NanoTag Biotechnologies | Fluorophore-conjugated NbPSD95 | IF, direct | 1:500 |
| Fig. 4 | | FluoTag-X2 anti-Syt1 AZDye568 | N2302-AF568-L | NanoTag Biotechnologies | Fluorophore-conjugated anti-Synaptotagmin1 nanobody | IF, direct | 1:500 |
| Fig. 4 | | FluoTag-X2 anti-GFAP Atto488 | N3802-At488-L | NanoTag Biotechnologies | Fluorophore-conjugated anti-GFAP nanobody | IF, direct | 1:500 |
| Fig. 5 | | FluoTag-X2 anti-PSD95 Sulfo-Cy3 | N3702-SC3-L | NanoTag Biotechnologies | Fluorophore-conjugated NbPSD95 | IHC, direct | 1:500 |
| Fig. 6 | | FluoTag-X2 anti-PSD95 AbberiorStar635P | N3702-Ab635P-L | NanoTag Biotechnologies | Fluorophore-conjugated NbPSD95 | IHC, direct | 1:500 |
| Fig. 6 | | Recombinant anti-Synaptotagmin antibody | 105 008 | Synaptic Systems | Rabbit monoclonal recombinant antibody | IHC, 1$^{st}$ Ab | 2 µg/mL |
| Fig. 6 | | FluoTag-X2 anti-Rabbit IgG AbberiorStar580 | N2402-Ab580-L | NanoTag Biotechnologies | Fluorophore-conjugated anti-Rabbit IgG2 nanobody | IHC, 2$^{nd}$ Ab | 1:500 |
| Fig. 7 | | Recombinant anti-PSD95 Antibody (Minibody), Rabbit Fc fusion | N3783 | NanoTag Biotechnologies | NbPSD95 fused to rabbit IgG Fc domain | IHC-P, 1$^{st}$ Ab | 1 µg/mL |
| Fig. 7 | | Secondary anti-Rabbit Biotin | 111-065-144 | Jackson Immuno Research | Biotin-SP AffiniPure Goat Anti-Rabbit IgG | IHC-P, 2$^{nd}$ Ab | 5 µg/ml |
| Fig. 7 | | Avidin-Biotin Complex (ABC)-HRP Kit | PK-4000 | Vector Laboratories | ABC-HRP Kit | IHC-P, amplification | prepared according to manufacturer's instructions |

**Table S1: Antibodies (continued)**

| Figure | Panel(s) | Product | Product number | Provider | Description | Application | Dilution/ Conc. |
|---|---|---|---|---|---|---|---|
| Fig. S1A | | Recombinant anti-PSD95 Antibody (Minibody), Mouse IgG1 Fc fusion | N3782 | NanoTag Biotechnologies | NbPSD95 fused to mouse IgG1 Fc domain | WB, 1st Ab | 1:1000 |
| Fig. S1A | | sdAb anti-MsIgG1 HRP | N2005-HRP | NanoTag Biotechnologies | HRP-coupled anti-Mouse IgG1 nanobody | WB, 2nd Ab | 1:4000 |
| Fig. S1B | | Recombinant anti-PSD95 Antibody (Minibody), Mouse IgG2 Fc fusion | N3785 | NanoTag Biotechnologies | NbPSD95 fused to mouse IgG2 Fc domain | WB, 1st Ab | 1:1000 |
| Fig. S1B | | sdAb anti-MsIgG2 HRP | N2705-HRP | NanoTag Biotechnologies | HRP-coupled anti-Mouse IgG2 nanobody | WB, 2nd Ab | 1:4000 |
| Fig. S1C | | Recombinant anti-PSD95 Antibody (Minibody), Rabbit Fc fusion | N3783 | NanoTag Biotechnologies | NbPSD95 fused to rabbit IgG Fc domain | WB, 1st Ab | 1:1000 |
| Fig. S1C | | sdAb anti-RbIgG HRP | N2405-HRP | NanoTag Biotechnologies | HRP-coupled anti-Rabbit IgG nanobody | WB, 2nd Ab | 1:4000 |
| Fig. S2A, Fig. S2B | 1st rows | FluoTag-X2 anti-PSD95 AbberiorStar635P | N3702-Ab635P-L | NanoTag Biotechnologies | Fluorophore-conjugated NbPSD95 | IF, direct<br>IF, direct | 14 nM<br>1 nM |
| Fig. S2A, Fig. S2B | 2nd rows | Recombinant anti-PSD95 Antibody (Minibody), Mouse IgG2 Fc fusion | N3785 | NanoTag Biotechnologies | NbPSD95 fused to mouse IgG2 Fc domain | IF, 1st Ab<br>IF, 1st Ab | 14 nM<br>1 nM |
| Fig. S2A, Fig. S2B | 3rd rows | Monoclonal anti-PSD95 antibody | MABN68 | Millipore | Monoclonal antibody (Mouse IgG2) | IF, 1st Ab<br>IF, 1st Ab | 14 nM<br>1 nM |
| Fig. S2A, B | 2nd/3rd rows | FluoTag-X2 anti-Mouse IgG2 AbberiorStar635P | N2702-Ab635P | NanoTag Biotechnologies | Fluorophore-conjugated anti-Mouse IgG2 nanobody | IF. 2nd Ab | 1:500 |
| Fig. S2 | | FluoTag-X2 anti-Syt1 AZDye568 | N2302-AF568-L | NanoTag Biotechnologies | Fluorophore-conjugated anti-Synaptotagmin1 nanobody | IF, direct | 1:500 |
| Fig. S2 | | FluoTag-X2 anti-GFAP Atto488 | N3802-At488-L | NanoTag Biotechnologies | Fluorophore-conjugated anti-GFAP nanobody | IF, direct | 1:500 |

**IF**: Immunofluorescence; **IHC**: Immunohistochemistry; **IHC-P**: Immunohistochemistry on paraffin-embedded sections, **WB**: Western blot; **1st Ab:** Primary antibody; **2nd Ab**: Secondary antibody
